# No Evidence for a Single Oscillator Underlying Discrete Visual Percepts

**DOI:** 10.1101/2021.01.05.425131

**Authors:** Audrey Morrow, Jason Samaha

**Affiliations:** University of California, Santa Cruz

**Keywords:** Discrete sampling, Flash-lag effect, Fröhlich effect, Alpha oscillations, Visual illusion

## Abstract

Theories of perception based on discrete sampling posit that visual consciousness is reconstructed based on snapshot-like perceptual moments, as opposed to being updated continuously. According to a model proposed by Schneider (2018), discrete sampling can explain both the flash-lag and the Fröhlich illusion, whereby a lag in the conscious updating of a moving stimulus alters its perceived spatial location in comparison to a stationary stimulus. The alpha-band frequency, which is associated with phasic modulation of stimulus detection and the temporal resolution of perception, has been proposed to reflect the duration of perceptual moments. The goal of this study was to determine whether a single oscillator (e.g., alpha) is underlying the duration of perceptual moments, which would predict that the point of subjective equality (PSE) in the flash-lag and Fröhlich illusions are positively correlated across individuals. Although our displays induced robust flash-lag and Fröhlich effects, virtually zero correlation was seen between the PSE in the two illusions, indicating that the illusion magnitudes are unrelated across observers. These findings suggest that, if discrete sampling theory is true, these illusory percepts either rely on different oscillatory frequencies or not on oscillations at all. Alternatively, discrete sampling may not be the mechanism underlying these two motion illusions or our methods were ill-suited to test the theory.

## Introduction

Whether conscious visual perception unfolds continuously or as a series of discrete updates remains a topic of much debate (Anliker, 1963; Doerig et al., 2019; Fekete et al., 2018; Harter, 1967; Herzog et al., 2016; James, 1890; Kristofferson, 1967a; VanRullen, 2016; VanRullen & Koch, 2003; Varela et al., 1981; C. T. White, 1963; P. A. White, 2018). Vision appears continuous to introspection, yet a number of perceptual phenomena are difficult to reconcile with a continuously updating perceptual process (Herzog et al., 2020; Sokoliuk & VanRullen, 2019). The flash-lag effect and the Fröhlich effect are two such examples. In the flash-lag illusion, a brief stationary stimulus (the ‘flash’) is misperceived as lagging behind a moving stimulus when the two stimuli are, in fact, spatially aligned (Metzger, 1932; Murakami, 2001; Nijhawan, 1994). The Fröhlich illusion refers to the observation that the onset of a moving stimulus is often mislocalized as being further along the trajectory of motion than it really is (Fröhlich, 1923; Kerzel, 2010). Many variants of discrete sampling models have been proposed (for reviews, see: Herzog et al., 2020; VanRullen, 2016; P. A. White, 2018), but recently Schneider (2018) proposed a unifying account of both the flash-lag and Fröhlich illusions based on discrete sampling.

In the model, a repeating process of sampling followed by reconstruction occurs, with the sampling process lasting for a specified duration (a ‘perceptual moment’). At the end of the perceptual moment, stimuli are registered in their last-known positions and this estimate forms the basis of the conscious reconstruction of events. Because moving stimuli will be in a different position at the end of the moment than a stationary stimulus, this leads to the kind of discrepancy observed between the flash and motion stimulus (flash-lag) or between the onset of the motion and the position the stimulus is in when conscious perception was updated (Fröhlich). Thus, the duration of an individual’s perceptual moment is the sole parameter in the model and half of this quantity corresponds to the average flash-lag and Fröhlich magnitude. This single-parameter model (Schneider, 2018) provided good fits to a large flash-lag dataset from Murakami (2001).

At the neural level, oscillations in brain activity have long been speculated to be involved in discrete perceptual sampling. For instance, within- and between-subject variation in alpha-band frequency (7-14 Hz) is predictive of temporal properties of visual (Baumgarten et al., 2018; S. Coffin & Ganz, 1977; Stephen Coffin, 1977; Gray & Emmanouil, 2020; Gulbinaite et al., 2017; Kristofferson, 1967b; Minami & Amano, 2017; Ro, 2019; Samaha & Postle, 2015; Shen et al., 2019) and cross-modal perception (Cecere et al., 2015; Cooke et al., 2019; Keil & Senkowski, 2017), with higher-frequency oscillations being associated with finer-grained temporal resolution. Moreover, the phase of ongoing alpha activity predicts perception of near-threshold visual stimuli (Alexander et al., 2020; Busch et al., 2009; Dugué et al., 2011; Mathewson et al., 2009; Samaha et al., 2015, 2017; Sherman et al., 2016). Indeed, it has been shown that trial-to-trial variability in the magnitude of the flash-lag effect is predictable by the phase of ∼7 Hz oscillations prior to flash onset (Chakravarthi & VanRullen, 2012), consistent with the model proposed by Schneider (2018). And a recent experiment presented a luminance-modulating annulus at 10 Hz surrounding a flash-lag display and found that the flash-lag magnitude was correlated with the phase of the luminance modulation, suggesting that entraining brain activity at an alpha frequency causes changes in flash-lag perception at that frequency (Chota & VanRullen, 2019).

The goal of this study was to test the proposition that, if the flash-lag and Fröhlich effects are driven by discrete sampling at the alpha frequency, then the magnitude of the illusion should be correlated across individuals. That is, given the relative stability of an individual’s alpha frequency (Grandy et al., 2013; Haegens et al., 2014), an individual with a large flash-lag magnitude (putatively caused by a lower alpha frequency and thus less frequent updating) should also have a large Fröhlich effect. Alternatively, no correlation between illusion strengths could indicate either that the Fröhlich and flash-lag do not reflect the same discrete sampling mechanism, or that they do reflect discrete sampling but at different frequencies that are not meaningfully related to one another across individuals. This latter notion is supported by a general lack of convergence in the literature on a single time scale of the ‘perceptual moment’ across tasks and stimuli, and a lack of a single oscillation frequency being related to temporal perception (Herzog et al., 2020; Ronconi et al., 2017; VanRullen, 2016). Thus, a correlation between illusion magnitudes is a necessary (but not sufficient) pre-condition for the theory that alpha sampling underlies both illusions. However, apart from playing a role in theorizing about discrete versus conscious perception, there is also a more general question of whether the flash-lag and Fröhlich effect are based on the same mechanism (whatever that may be), which remains controversial (Kreegipuu & Allik, 2003; Krekelberg & Lappe, 2000; D. Whitney & Cavanagh, 2000) but which can be informed by an individual differences approach.

We quantified the magnitude of the flash-lag and Fröhlich illusion psychophysically by determining the spatial offset required between a motion stimulus and a stationary reference stimulus to make the two stimuli appear spatially aligned (i.e., the point of subjective equality; PSE). Robust flash-lag and Fröhlich illusions were present in our displays and despite the two displays being highly similar, we observed no correlation between individual differences in the illusion magnitudes. A Bayesian analysis provided moderate support for the null hypothesis of no correlation. We conclude that either 1) our stimulus parameters were ill-suited to detect a true relationship, 2) these two illusions are supported by different mechanisms, perhaps based on sampling rates determined by different oscillatory frequencies in the brain or 3) distinct mechanisms not based on discrete sampling underlie each illusion.

## Method

### Participants

Twenty-four participants (8 male, 1 prefer not to say; age range: 18-29) were recruited from the University of California Santa Cruz’s (UCSC) online psychology research pool. The experiment was approved by the UCSC Institutional Review Board. All participants had normal or corrected-to-normal vision and provided written consent to participate. One participant’s data was excluded for indicating, during debriefing, that they did not understand the response mapping. As a result, their data did not follow a typical psychometric curve.

A posthoc power analysis (G*power 3.1; Faul et al., 2007) indicated that 23 participants achieved 80% power to detect a one-tailed correlation of magnitude 0.5 (alpha level = 0.05). Given that the theory we are testing proposes that both illusions are based on the same mechanisms of alpha-band sampling, this would predict a strong correlation between the two illusion magnitudes. Our group has published similarly sized samples when investigating individual differences in theoretically-large associations (Samaha & Postle, 2015, 2017). However, we sought to quantify more precisely the degree of evidence in favor of the null hypothesis provided by our results by using Bayesian analyses (described later).

## Data Availability

In accordance with the practices of open science and reproducibility, all raw data and analysis scripts will be made publicly available on the Samaha Lab’s Open Science Framework repository (https://osf.io/ahm6c/) upon publication of this manuscript.

### Design

The experiment took place in a dimly lit room with participants positioned in a headrest to maintain a distance of 74cm from the computer monitor. Stimuli were generated using Psychtoolbox-3 (Kleiner et al., 2007) running in the MATLAB environment (version 9.8) under the Ubuntu (version 18.04) operating system. Stimuli were presented on a middle-gray background of 50cd/m^2^ luminance on a gamma-corrected VIEWPixx EEG monitor (1920 x 1200 pixels, 120 Hz refresh rate). A central fixation, consisting of a gray crosshair superimposed over a black circle 0.4° of visual angle, was present on the screen throughout each task.

#### Flash-lag Illusion

A 0.3° black dot moved horizontally outwards from the center in either the left or right hemifield (randomly determined on each trial with equal probability). The dot trajectory started at 2° from central fixation and moved horizontally away from fixation for 2000ms at a speed of 6° per second. At a randomly selected interval between 500-1500ms during the dot’s trajectory, a set of vertical black bars (the “flash”) flashed above and below the dot for 8ms. The two bars were 0.13° W x 0.83° H with a distance of 0.83° between them. The flash appeared at one of 8 offsets (randomly selected with equal probability on each trial) from the set [-0.70°, −0.40°, − 0.24°, −0.08°, +0.08°, +0.24°, +0.40° +0.70°] relative to the location of the moving dot, which translates to between 4.3° to 11.7° away from central fixation. This variation in offset caused the flash to appear “ahead of” or “behind” the dot to varying extents (see Figure 1).

**Figure 1.**
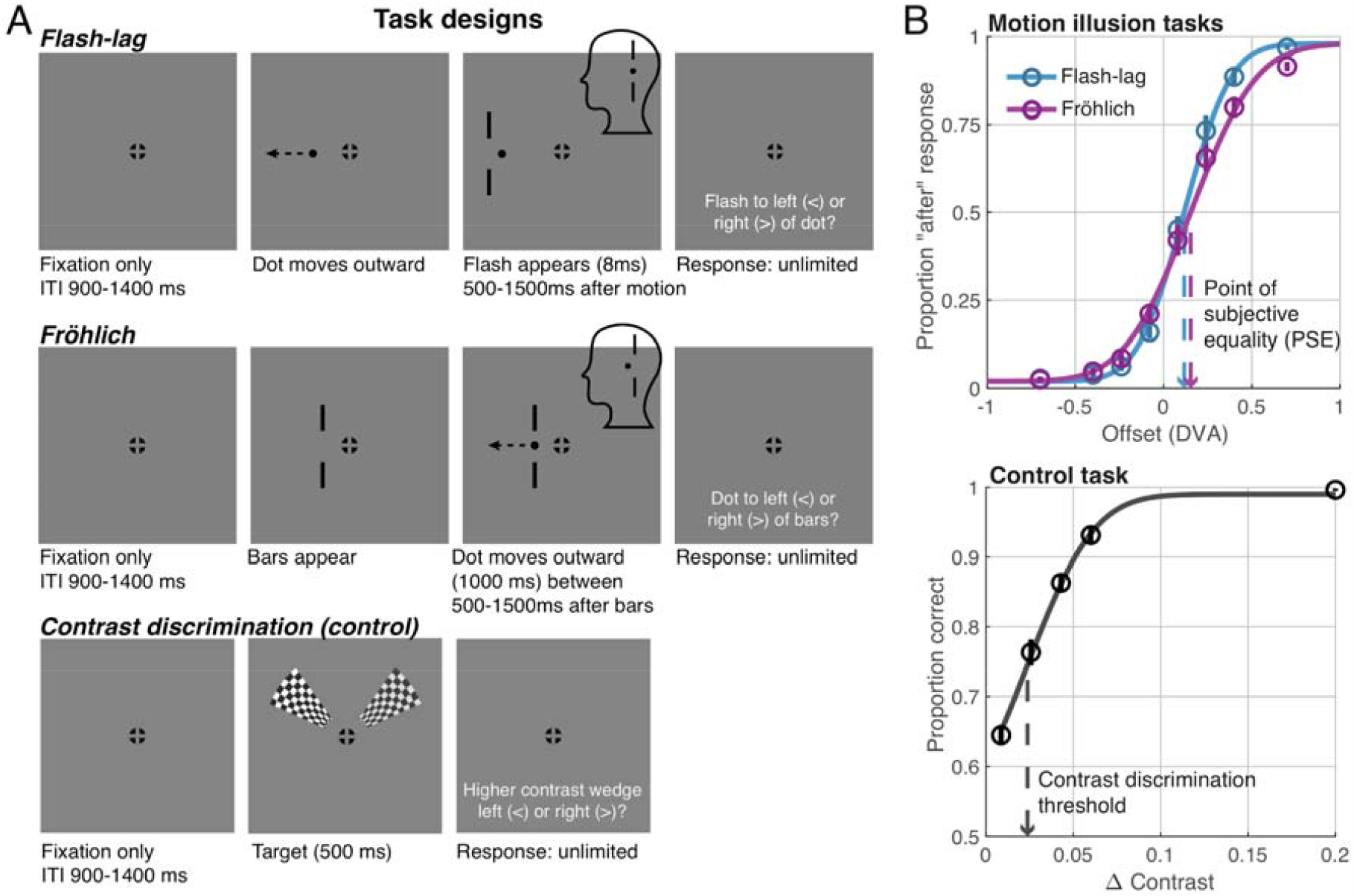
Task diagrams and group-level psychometric functions. **A)** From top-to-bottom, example trails of the flash-lag, Fröhlich, and control tasks are shown. For the two illusion tasks (flash-lag and Fröhlich, respectively), participants judged the location of the vertical bars relative to the moving dot, or the initial location of the moving dot relative to the bars. All that differed between tasks was whether the moving dot appeared before the vertical bars (flash-lag condition) or after (Fröhlich) and whether the bars were presented briefly (8ms; flash-lag) or remained on screen throughout the trial (Fröhlich). In the example flash-lag trial, a positive offset is shown which refers to the fact that the vertical bars flashed further away from fixation than the moving dot. The cartoon head insert represents the perceived illusion and was not part of the display participants saw. In the flash-lag effect, the flashed bars appear to lag behind the motion stimulus which, for a positive offset, would manifest as the bars appearing to be aligned with the dot. In the example Fröhlich trial, the offset equals zero, but the initial position of the dot is perceived as advanced along the trajectory of motion. In the contrast discrimination (control) task, two wedges were shown simultaneously, one with higher contrast (the left wedge in this example) and participants judged the location with highest contrast. The white text shown in these schematics was not present on the actual displays. **B)** Group-level behavior (circles) and psychometric function fits (lines) for the two illusion tasks (top) and control task (bottom). In both the flash-lag and Fröhlich data, the PSE is reliably non-zero, indicating that a positive offset between the dot and bars are required for participants to perceive the two stimuli as being aligned. Contrast discrimination thresholds provided control data in that they were hypothesized to be independent of the illusion PSEs. Error bars represent ±1 SEM across subjects and sometimes too small to be clearly visible.

#### Fröhlich Illusion

This task used the same parameters as the flash-lag for the moving dot and the set of bars except that the bars appeared first (either to the left or right if fixation, randomly determined), and remained on screen for 2000ms. The dot then appeared midway into the 2000ms (between 500-1500ms) and immediately began moving (see Figure 1). The bars were placed using the same set of possible offsets between the onset of the dot trajectory ([-0.70°, −0.40°, −0.24°, − 0.08°, +0.08°, +0.24°, +0.40° +0.70°], randomly selected on each trial), which translates to between 5° to 11° from fixation. Similar to the flash-lag illusion, this offset caused the dot to appear “ahead of” or “behind” the set of bars to varying degrees.

#### Contrast Discrimination (Control) Task

In addition to the two illusion tasks, we included a third control task that was intended to measure perceptual processes (here, contrast discrimination) clearly different from those underlying the flash-lag and Fröhlich illusions. The control task was adapted using stimuli from Iemi et al. (2019) and presented two checkerboard wedges to the left and right of fixation (see Figure 1). The wedges corresponded to segments of an annular checkerboard with a spatial frequency of 5 cycles per degree. The two wedges appeared for 500ms in either the upper or lower hemifield with the inner edge of the wedges 3° from central fixation and the outer edge 10° away. The right wedge was designated a ‘standard’ and was held constant at 0.8 contrast (i.e., 80% Michelson contrast) and the left wedge varied randomly across the following 10 levels [0.60, 0.740, 0.757, 0.774, 0.791, 0.808, 0.825, 0.842, 0.860, 1.0]. This task was a control in the sense that the specificity of any correlation observed between the flash-lag and Fröhlich illusions could be tested by comparing this effect to the correlation obtained between the illusion PSEs and contrast discrimination thresholds (which, we hypothesized, would be uncorrelated).

### Procedure

Participants were instructed on each task before engaging in practice blocks of 20 trials. Researchers monitored the practice blocks to ensure participants understood the task instructions and reviewed the response curves after each block. Additional practice blocks were conducted as necessary. The order of the three tasks was counterbalanced across participants. For all tasks, the intertrial interval varied randomly between 900ms and 1400ms and response duration was unlimited so that participants would focus on accuracy over speed of response. Participants were instructed to maintain fixation on the central crosshair throughout the whole trial and not to track the moving object.

Participants completed 312 trials, split into three blocks, of each illusion task. Participants were instructed to respond using the “<“and “>“keys to indicate whether the new object appeared to the left or right of the original object, respectively. In other words, participants responded to whether the flash appeared to the left or right of the moving dot in the flash-lag illusion or to whether the moving dot first appeared to the left or right of the stationary bars in the Fröhlich illusion.

Participants completed 300 trials of the control task, split into three blocks. In this task, participants engaged in contrast discrimination between the two checkerboard wedges. The participants responded with the same “<“and “>“keys to indicate whether the left or right wedge, respectively, had a greater level of contrast.

### Data Analysis

#### Psychometric functions

Prior to fitting, data from the right and left hemifield of the illusion tasks were collapsed and responses were mirrored to remove relative location information (i.e responses were recoded to reflect “before’’ or “after” responses rather than “left” or “right” responses, the meaning of which was dependent on the hemifield of presentation). When analyzing data from the contrast discrimination task, the contrast of the variable-contrast wedge was expressed in units of absolute difference from the standard contrast wedge (i.e., □ contrasts = [0.008, 0.025, 0.042, 0.060, 0.2]) and accuracy (proportion correct) was computed for each □ contrast level.

A cumulative normal function was fit to each subject and task using maximum-likelihood estimation as implemented in the Palamedes toolbox (version 1.9.1; Prins & Kingdom, 2018). For the flash-lag and Fröhlich task, the PSE and slope were free parameters whereas the lower and upper asymptotes of the curve were fixed at 0.02 and 0.98, respectively, to allow for lapses (Prins, 2012). For the contrast discrimination data, the threshold and slope were free parameters and the guess rate was fixed at 0.5 and lapse rate at 0.02. The across-subject correlation between psychometric parameters in each task was then computed using a Spearman correlation (rho), to mitigate the influence of any potential outliers.

#### Bayes factor analysis

To interpret the weight of evidence our data provide for and against the null hypothesis of no correlation, we computed Bayes factors (BF). Here, BF reflects the ratio of likelihoods (L) of the data under the alternative hypothesis (the theory; H1) to that of the null hypothesis (H0); that is BF_1,0_= L_H1_/L_H0_. Thus, a BF increasing from 1 indicates more evidence for H1, whereas a BF approaching zero indicates increasing evidence for H0. Meaningful interpretation of a BF requires specifying an appropriate H1, which amounts to specifying a theoretically-plausible distribution of effects according to a theory (Dienes, 2014; Rouder et al., 2016). We specified two models of H1, corresponding respectively to weak and strong versions of the theory that alpha-based sampling underlies both illusions.

The first model instantiates the relatively minimal assumption that the theory just predicts a positive correlation (rho) between the flash-lag and Fröhlich effect. This was specified in the Matlab implementation of the BF calculator by Zoltan Dienes (http://www.lifesci.sussex.ac.uk/home/Zoltan_Dienes/inference/Bayes.htm) as a uniform distribution spanning 0 (lower bound of rho) to 0.9 (upper bound of rho; values greater than 0.9 were considered unrealistic). (Note that values were Fisher’s z-transformed prior to input into the calculator.) This expresses the view that the theory predicts some positive relationship that is equally likely to be of small, medium, or large effect size. We refer to this as BF_U(0, 0.9),0_. The actual mean and spread of the observed effect (“sample mean” and “sample SE” in the calculator) were taken as the Fisher’s z-transform of the actual rho value computed between the illusions with SE = 1/sqrt(df - 1).

The second BF represents H1 in a more theoretically-motivated way using the ratio-of-scales heuristic from Dienes (2019). This approach rests on the logic that, if the two illusions are caused by the same underlying mechanism, and because both are measured in the same units (PSE in degrees of visual angle, or DVA), a strong version of the theory predicts that they should be identical. That is, the slope (beta) of a line fit to the Fröhlich by flash-lag PSEs (e.g., Figure 3A) should equal 1. This predicted effect size for H1 was specified in the Dienes code as a half-normal distribution (to reflect the directional nature of the hypothesis) with a mean of zero and an SD of beta/2 = 0.5. Since the prediction that beta = 1 is a maximum effect that assumes no error, the specification of a half normal with a mean of zero is a conservative estimate of the H1 mean since it predicts that smaller effects are more likely. Dividing the predicted value by 2 to achieve an SD of 0.5 is a further conservative correction that halves the predicted effect, as recommended in the literature (Dienes, 2014, 2019). We refer to the BF according to this model of H1 as BF_HN(0, 0.5),0_. The observed beta and SE from the data were estimated as the slope and standard error of an ordinary least-squares regression predicting the flash-lag PSE from the Fröhlich PSE and were input into the BF calculator.

## Results

We quantified the magnitude of the flash-lag and the Fröhlich illusion within the same individuals to assess their correlation. The data were well-described by a psychometric function (R^2^ across all subjects in both tasks: mean = 0.98, range = [0.92, 0.99]; see Figures 1 and 2). As a first step, we tested whether illusory percepts consistent with the flash-lag and the Fröhlich effects were observed in our displays. Because each illusion was quantified as the PSE whereby the offset between the dot and bar stimuli would be judged as either “before” or “after” 50% of the time, a PSE of zero DVA would correspond to veridical perception. Consistent with illusory percepts, however, the mean (±SEM) PSE in the flash-lag task was 0.12 (±0.02) DVA, which was significantly different from zero (t(22) = 5.74, p < 0.0001) and the mean Fröhlich PSE was 0.15 (±0.03) DVA, which also differed from zero (t(22) = 5.43, p < 0.0001). A positive PSE indicates, in the flash-lag case, that the bars needed to be flashed further from fixation than the moving dot by 0.12 DVA in order to be perceived as aligned with the dot (i.e., to offset the illusory lag of the flash). In the Fröhlich task, the bars needed to be 0.15 DVA further from fixation than the onset of motion in order for the motion onset to be perceived as aligned with the bars (consistent with a misperception of the motion stimulus as being advanced along the motion trajectory). The two illusions were of a comparable magnitude as they did not significantly differ from one another (t(22) = 1.05, p=0.304). Under comparable conditions to those used here, similar magnitude illusions have been reported for both the flash-lag and Fröhlich effect (Eagleman & Sejnowski, 2007). Contrast thresholds from the control task were, on average, Δ0.024 (±0.002), indicating that observers could discriminate, on average, a contrast difference between wedges of 2.4% with 75% accuracy.

**Figure 2.**
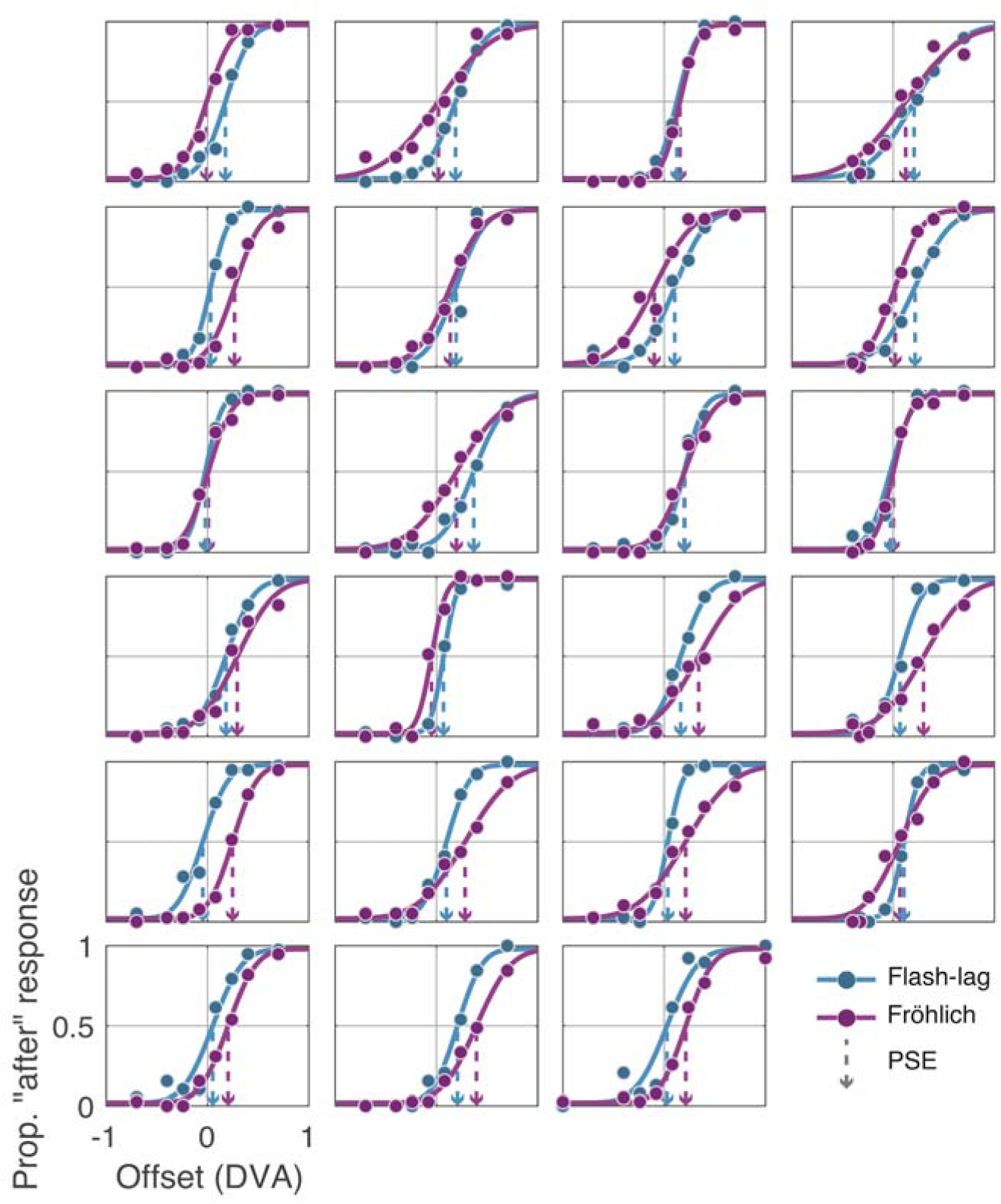
Individual subject data (circles) and psychometric function fits (lines) for the two illusion tasks. Dashed lines denote the PSE for each subject.

We next asked whether the magnitude of an individual’s flash-lag illusion (flash-lag PSE), was predictive of the magnitude of their Fröhlich illusion (Fröhlich PSE), as would be expected if the two illusions result from the same underlying oscillatory sampling frequency. However, the across-task correlation in PSEs was virtually zero (rho = −0.0089, p = 0.969, 95% bootstrap CI = [-0.41, 0.39], indicating no evidence for a relationship between these illusions (Figure 3A). As evident from the bootstrap analysis presented in Figure 3A, the true correlation could plausibly span a wide range between ±0.4, though the mean of this bootstrap distribution is virtually zero (−0.006). A Bayesian analysis (Dienes, 2014) that quantified the likelihood of the data belonging to the null hypothesis (H0) of no correlation versus the alternative hypothesis (H1) of a positive correlation (a weak version of the theory), indicated that the data are 5.56 times more likely to have been generated under the null (BF_U(0, 0.9),0_ = 0.179). Using a hard version of the theory to model H1, according to which the two PSEs are equal, resulted in a BF_HN(0, 0.5),0_ = 0.419, indicating that the null is 2.38 times more likely than the alternative. Thus, depending on the exact implementation of the theory, the data are approximately 2 to 5 times more likely under the null.

**Figure 3.**
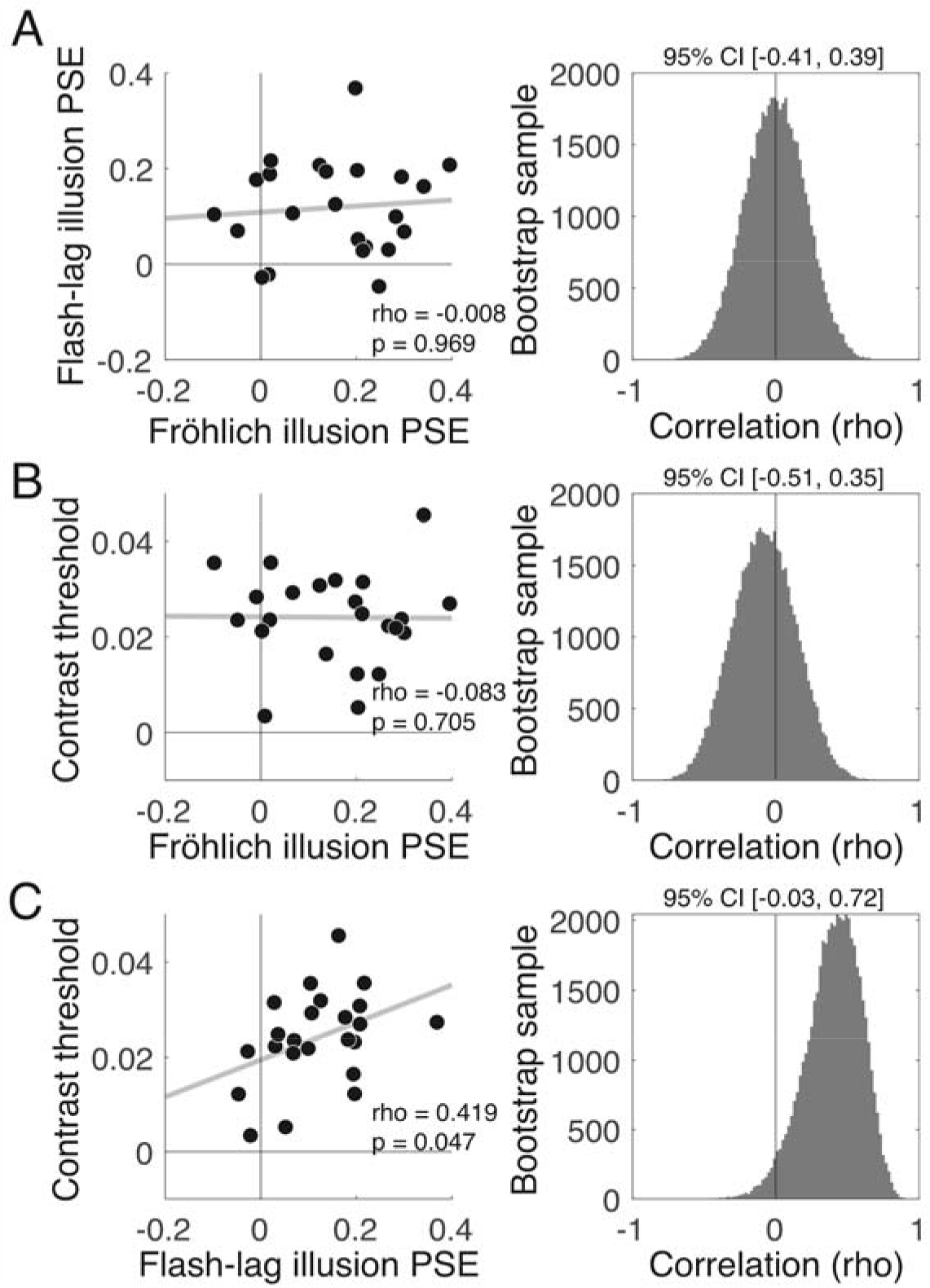
Across task correlations between psychometric thresholds. **A)** Correlation (left) and bootstrap analyses (right) revealed virtually zero correlation between the flash-lag and Fröhlich illusion PSE. This indicates that the magnitude of one illusion is not predictive of the magnitude of the other. **B)** No correlation was found between Fröhlich PSE and contrast thresholds. **C)** An unexpected medium-sized positive correlation was observed between flash-lag PSE and contrast threshold, indicating that an individual capable of discriminating a small change in contrast has a smaller magnitude flash-lag effect. We speculate that this may be due to the very brief flash used in our display (8ms), accurate perception of which may be aided by a lower contrast threshold. Lines of best fit are shown in grey and Spearman rho values with 95% bootstrap CI are provided.

Next, we tested for associations between contrast thresholds and PSEs in each illusion task. As shown in Figure 3C, contrast thresholds were positively correlated with flash-lag PSEs (rho = 0.419, p = 0.0478, 95% bootstrap CI = [-0.03, 0.72]), indicating that individuals capable of discriminating smaller differences in contrast had smaller flash-lag illusions. (We speculate on the reason behind this correlation in the discussion.) No correlation was observed between contrast thresholds and the Fröhlich PSE (rho = −0.083, p = 0.706, 95% bootstrap CI = [-0.51, 0.35]). Because contrast thresholds unexpectedly explained variance in the flash-lag effect, we sought to test if an effect of Fröhlich PSE on flash-lag PSE emerged when controlling for the influence of contrast discrimination. However, a multiple regression model that predicted individual differences in the flash-lag PSE using the Fröhlich PSE and contrast thresholds (as a covariate) did not reveal an association between the two illusions (beta = 0.066, SE = 0.148, t = 0.447, p = 0.659), consistent with the results from the main correlation analysis.

## Discussion

This study assessed whether the same mechanism, discrete perceptual sampling at a single frequency, could explain the flash-lag and Fröhlich illusions. We robustly induced illusory percepts in both of the flash-lag and Fröhlich displays, yet the correlation between illusion sizes was virtually zero. A Bayesian analysis allowed us to determine the likelihood of our data being obtained from the hypothesized distribution according to alpha-based sampling. The BF indicated that responses to the flash-lag and Fröhlich illusion tasks were between 2.4-5.5 times more likely to come from the null distribution than various theory-derived alternatives, suggesting it is unlikely that a single oscillator is underlying individual variations in the flash-lag and Fröhlich effects.

Although we observed moderate evidence for a null effect, there are several underlying reasons for obtaining null results. It could still be that discrete sampling theory is true and is neurally instantiated via oscillations, but that these two illusions rely on different frequencies which are themselves uncorrelated. A large body of work has linked alpha-band oscillations to temporal windows of processing (Cecere et al., 2015; S. Coffin & Ganz, 1977; Cooke et al., 2019; Grabot et al., 2017; Gray & Emmanouil, 2020; Kristofferson, 1967b; Minami & Amano, 2017; Samaha & Postle, 2015; Shen et al., 2019; Varela et al., 1981; Wutz et al., 2018) and even specifically to the flash-lag illusion (Chakravarthi & VanRullen, 2012; Chota & VanRullen, 2019). On the other hand, lower-frequency oscillations in the theta range are also often implicated in establishing temporal windows of perception (Nakayama et al., 2018; Wutz et al., 2016; for review see VanRullen, 2016). Indeed, even different tasks within the same subjects can reveal different frequencies related to the temporal parsing of visual stimuli (Ronconi et al., 2017), indicating that multiple and task-dependent rhythms may underlie the perceptual moment. Thus, our results are compatible with a model according to which the flash-lag and Fröhlich effect are driven by discrete sampling but at different frequencies. However, on this theory, our finding that the two illusions were of very similar magnitude (Figure 1) may still require further explanation.

A second possible interpretation of our null effect is that these illusions do not rely on discrete sampling at all, contra Schneider (2018). Other viable accounts of the flash-lag and Fröhlich effects have been put forth and recently defended. Regarding the flash-lag, a recent review argues that motion extrapolation is currently the best account (Hogendoorn, 2020). Motion extrapolation refers to the prediction of a moving object’s trajectory based on its recent past. This is distinct from discrete sampling (or the related post diction account of the flash-lag) in that the motion percept is not reconstructed after the fact, but is instead based on a prediction about where the stimulus will be. Motion extrapolation mechanisms have been observed as early as the retina (in some species) and could therefore begin very early after motion onset to produce a percept of the motion stimulus that is advanced with respect to a stationary flash (Hogendoorn, 2020). Regarding the Fröhlich effect, it has been argued that metacontrast masking plays a crucial role (Kerzel, 2010). In metacontrast masking, the mask and target do not overlap and stimulus onset asynchronies between 40-100ms between target presentation and mask presentation create strongest masking effects (Kerzel, 2010). It has been proposed that metacontrast masking suppresses the initial trajectory of the moving stimulus, thus creating the illusion that it begins farther ahead than its true starting location (Piéron, 1935). Because the moving dot is already along its trajectory when the flash appears in the flash-lag effect, this masking effect of the beginning trajectory would not affect perception in the flash-lag illusion. Thus, it remains an open possibility that these two illusions are based on different mechanisms, either one of which may not be discrete sampling.

A third possibility is that our stimulus design was suboptimal for detecting a true relationship. For instance, an unexpected correlation was found between PSEs in the flash-lag illusion and the contrast discrimination task. The control task was primarily administered to rule out the possibility that any observed correlation between the two illusion PSEs was a trivial reflection of *any* two tasks being correlated across individuals. Although this point is moot since no correlation was observed between the two illusion tasks, we speculate that the observed correlation between contrast thresholds and the flash-lag PSE might have occurred since the flash in our flash-lag display was very brief (8ms) and perhaps having a lower contrast threshold could translate to more veridical flash localization and a smaller flash-lag effect. However, a multiple regression model of our data that included contrast discrimination thresholds as a covariate still failed to find any relationship between the two illusions. A longer flash should reduce variance in flash-lag estimates due to individual differences in contrast perception. Future studies should consider increasing the flash duration to facilitate perception of the flash, while also keeping in mind how the speed and proximity of the moving dot may interact with a longer flash and impact illusory effects. An additional stimulus consideration that may affect both the flash-lag and the Fröhlich effect is the interaction of each illusion with low-level motion processing. The flash-drag effect suggests that moving stimuli may distort the surrounding visual space and, thus, would affect perception of any nearby stationary stimuli (Whitney & Cavanagh, 2000). Moving the bars farther away from the horizontal trajectory of the dot in both illusion tasks could help to mitigate any distortion caused by low-level motion processing.

Although it can be difficult to know the source of a null effect, even if evidence in support of the null can be shown (as with our Bayesian analysis) our data can inform future studies. The task parameters used in this study seem to effectively produce the expected illusory effects of the flash-lag and Fröhlich illusions and such parameters could be used to test other perceptual phenomena or mechanisms related to discrete sampling theory. Alternatively, adjusting these task parameters or comparing the PSE across additional tasks could provide further insight into how individuals sample visual information, or more specifically, what stimulus designs might better capture the hypothesized ‘perceptual moments.’ Overall, our current data do not support the hypothesis that individual alpha frequency might be the mechanism underlying discrete sampling theory, but these findings are not sufficient to rule out discrete sampling theory altogether.

## Acknowledgements

The authors would like to thank Leo Whisten for help with data collection.

